# Unravelling Antibody-Induced Mechanical Stability of Antigen: Insights from Single-Molecule Studies

**DOI:** 10.1101/2024.05.24.595683

**Authors:** Soham Chakraborty, Shivam Pandit, Madhu Bhatt, Debojyoti Chowdhury, Suman Chakrabarty, Shubhasis Haldar

## Abstract

The intricate nature of antigen-antibody interactions plays a pivotal role in immunological responses. Despite the multitude of ligand-binding sites on antigens, the influence of antibodies on their mechanical stability remains elusive. This study elucidates the impact of IgM, the largest antibody isotype, on the mechanical stability of protein L, a bacterial superantigen, using single-molecule magnetic tweezers and steered molecular dynamics. Our findings reveal a concentration-dependent elevation in mechanical stability induced by IgM, as demonstrated by prolonged unfolding dwell times. Through steered molecular dynamics simulations, we elucidate the distinct mechanical responses of protein L binding interfaces at various IgM complex states, highlighting their synergistic effect on IgM dimer complex stability. Notably, this enhanced response stems from the altered unfolding pathway of protein L upon IgM interaction, providing significant insights into the generic mechanisms governing antibody-induced mechanical stability of antigenic substrates in physiological conditions, shedding light on the underlying folding dynamics and molecular mechanics of antigen-antibody interaction.

## Introduction

Antibody interaction can impact the antigen stability and therefore their function through the binding interfaces. Specific interactions between antibody determinant (paratope) and antigenic determinant (epitope) are physical in nature through different short and long-range attractive forces such as electrostatic and hydrophobic interactions, which occur at several nanometer range^1^. From a physical perspective, these biomolecular interactions can change their affinity by several orders of magnitude under different force constraints and can change the substrate mechanical stability. Atomic force microscopy have already been implemented to study the adhesion force between such protein-pairs including biotin-avidin pair and antigen-antibody complexes^2–4^. It has also been reported that small ions as potent ligands modulate the protein stability without significant structural perturbation^5,6^. These effects are generally predictive owing to the presence of single binding sites in the receptors. However, the stabilizing effect of ligand could be different in the case of antigens containing more than one antibody-binding interfaces. Additionally, antibody-induced stability of antigens promote their ability to function under mechanical sheer of mucus flow or urination^5^. Since the ligand-induced mechanical stability of protein is connected to their conformational trajectory, a clear understanding of the structural changes of antigen upon antibody engagement is required. However, a detailed understanding of how antibody molecules increase the mechanical stability through the binding interfaces of proteins under the mechanical sheer is poorly understood in the literature.

To address these queries, we explored the mechanical impact of the largest antibody isotype IgM on superantigen protein L B1 domain mechanical stability by magnetic tweezers, implementing its force-clamp methodology that can apply a constant force on a substrate protein^7^. Moreover, the benefit of a broader force range spanning from 0 to 120 pN, coupled with precise force resolution, enables us to investigate the influence of antibodies on protein stability. Finally, using this instrumental set-up, the force can be precisely applied on the substrate protein while keeping the antibody molecules unperturbed.

We systematically investigated the impact of IgM antibodies on the mechanical stability of protein L, a model super-antigen found on bacterial surfaces known to interact with human antibodies^8,9^. Our magnetic tweezer experiments showed that antibody binding significantly increases the unfolding dwell time of protein L, thereby enhancing its mechanical stability in a concentration-dependent manner. To understand the molecular basis of this binding-induced stability, we performed steered molecular dynamics (SMD) simulations on protein L complexes, ranging from their apo state to IgM-bound states via different binding interfaces in protein L. Notably, we observed unique mechanical responses of each IgM-binding interface, which exert additive effect on the overall protein L stability under mechanical force. Our simulations indicate that the enhanced mechanical stability of protein L induced by antibody binding is due to conformational changes upon engagement with the antibody. Furthermore, our results suggest that the antibody-induced mechanical stability of protein L originates from conformationally-altered unfolding mechanism upon antibody engagement: while in apo-state, protein L unfolding initiates from the intrachain contacts breaking between its two parallel β-sheets, which triggers the unfolding of N-terminal β-sheets, followed by the loosening of intrachain contacts of C-terminal β-sheets with α-helix; in IgM-bound state, C-terminal β-sheets are more susceptible to unfold under mechanical constraints, followed by N-terminus. This intricate IgM-binding mechanism of protein L may regulate its mechanical stability in response to varying antibody concentrations during different phases of infection, thereby offering a fundamental understanding of antibody-induced mechanical responses in antigenic proteins under physiological shear stress.

## Results

### Magnetic tweezer study revealed a more mechanically stable IgM-bound protein L conformation

Protein L polyprotein construct, inserted between a C-terminal AviTag and N-terminal HaloTag has been used to perform the magnetic tweezers experiments^7,10,11^. While C-terminal end of the substrate is attached with a streptavidin-coated paramagnetic bead, the N-terminus is tethered to a glass surface via HaloTag chemistry^7,10^. The applied force by the permanent magnets is an inverse function of the distance between the paramagnetic bead and permanent magnet and characterized as an increase in the step size (defined as the difference between the folded and unfolded states) (Fig. 1A). The detailed force calibration method of our instrumental set-up has been previously reported^11,12^. Fig. 1B showed the representative traces of protein L obtained from the single-molecule experiment. At first, the polyprotein remains folded at 4 pN, which is then completely extended by an unfolding pulse of 75 pN that is visualized as stepwise increase of ∼15 nm in the polyprotein length. For native protein L (without IgM), it takes only ∼5 s to unfold all the domains, while increases to ∼58 s in the presence of 6 µM IgM (Fig. 1B). This increase in unfolding dwell time indicates that antibody interaction mechanically stabilizes the protein L under force.

**Figure 1:**
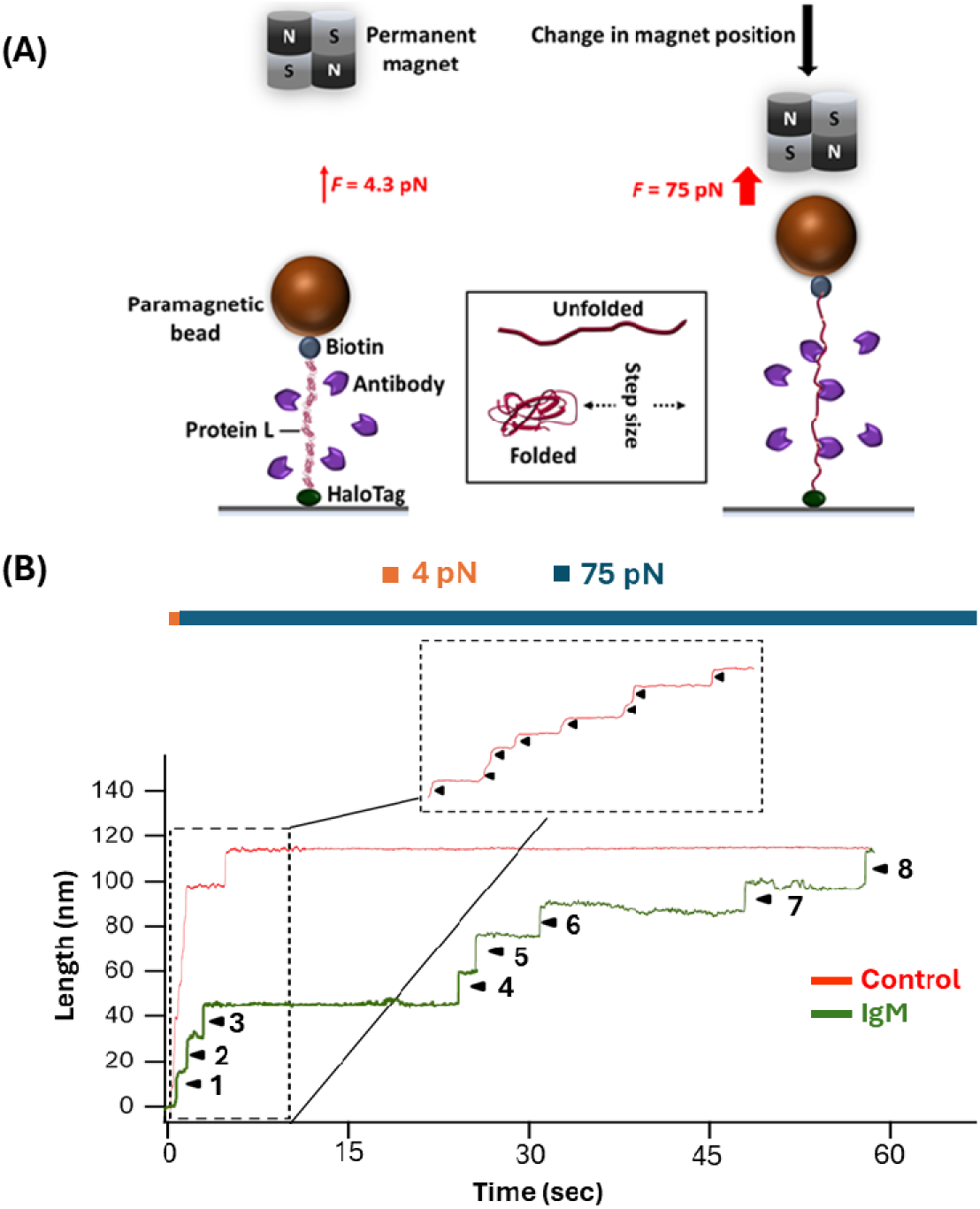
Single-molecule magnetic tweezers setup to study the protein L-antibody interaction: (A) A schematic illustration of magnetic tweezers experiment: In the polyprotein construct, tandem repeats of eight protein L domains are inserted within AviTag and HaloTag, helping them to attach with the paramagnetic bead through AviTag and to the glass surface via HaloTag. The applied force on the attached substrate can be controlled by regulating the distance between the permanent magnet and the paramagnetic bead attached to the Protein. As the substrate protein undergoes unfolding under tension, it extends, creating a step size defined by the difference between the lengths of the unfolded and folded domains. **(B) Exemplary unfolding profile of protein L during interaction with IgM:** At first, the protein L polyprotein remains folded at 4 pN, and applying a pulse of 75 pN unfolds all the eight domains of the construct. In the absence of IgM antibody (red), the polyprotein completely unfolds within ∼5 s; however, it increases to ∼58 s in the presence of 6 µM IgM (green), suggesting that IgM mechanically stabilize the protein L under force.

### Protein L contains two binding sites for the IgM interaction measured by probability histogram analysis

To gain in-depth understanding of how antibody affects the protein L unfolding kinetics under force, we have performed the force-clamp experiment of protein L at 75 pN force in the presence of different IgM concentration up to 6 µM. The unfolding dwell times of protein L has been plotted as probability distribution function (PDF) and fitted to the Gauss equation. The most probable unfolding dwell time of protein L at IgM-unbound state (control) has been observed to be 1.97 s (Fig. 2A). Interestingly, IgM increases this unfolding time steadily to 33.65 and 66.1 s at 1 µM and 6 µM concentration, respectively (Fig. 2B and 2C). To explore the unfolding events arising from antibody-bound Protein L at different concentration, we have plotted the most probable unfolding dwell time against different IgM concentration and fitted to a double exponential equation, indicating the presence of two binding sites for IgM antibodies (Fig. 2D). Furthermore, to investigate the IgM-induced mechanical stability of protein L, we have extended the polyprotein construct through several force-ramp cycles by increasing the force from 4 to 80 pN at different loading rates (1.2 to 4.3 pN/s), which have been suggested to be physiologically relevant^13–15^. In the force-ramp cycles, the unfolding steps are vertically extended to the force-extension curve to estimate the unfolding forces (Fig. 3A). We have plotted these unfolding forces of protein L at different unfolding rates, in the presence of IgM and have found to follow the Bell-Evans distribution. We observed that at zero-loading rate, the unfolding force of protein L is 32.98±0.93 pN in the absence of IgM which elevates to 60.54±1.63 pN in the presence of 6 µM IgM (Fig. 3B), signifying to increased mechanical stability upon IgM interaction.

**Figure 2:**
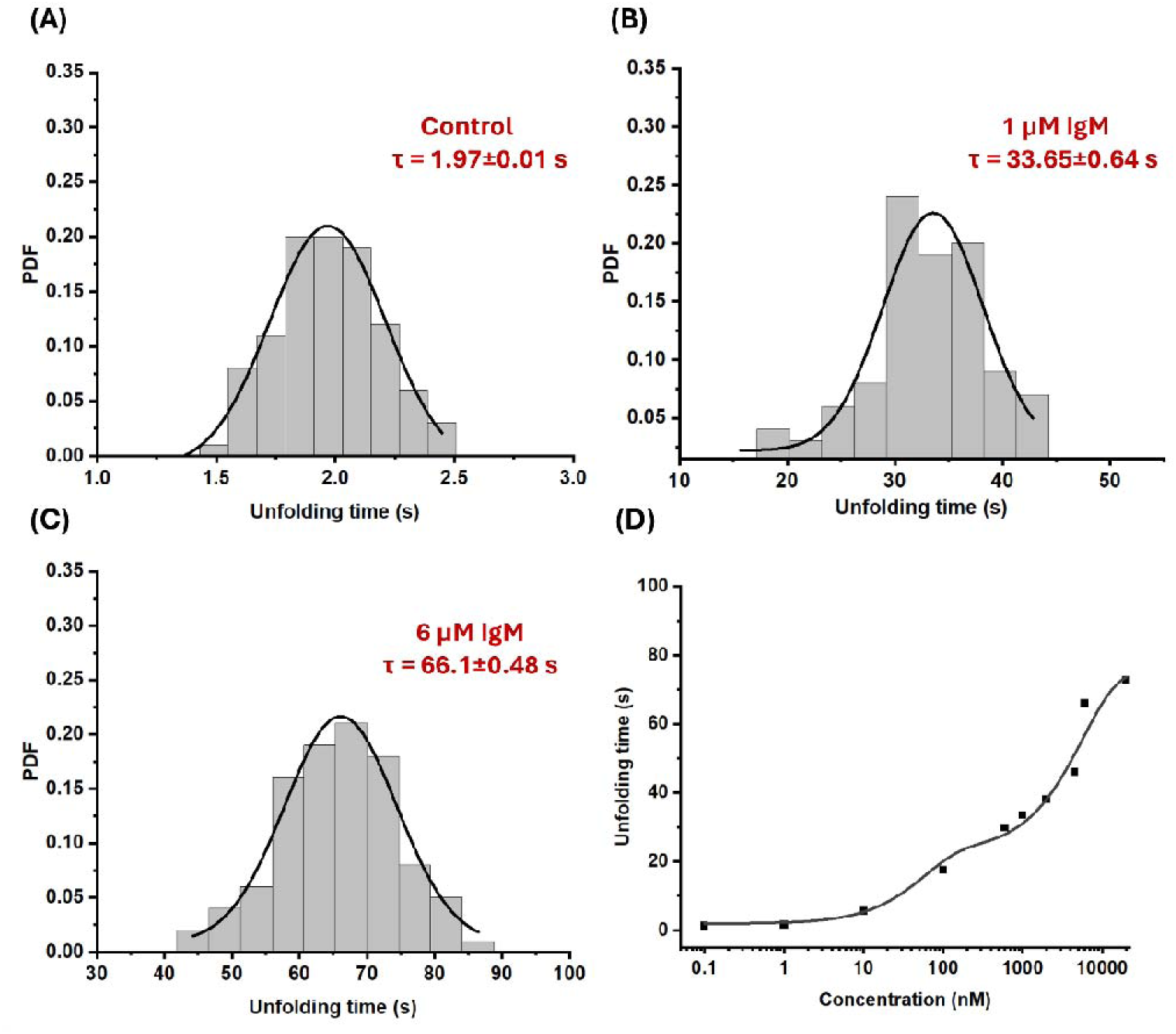
Unfolding kinetics of protein L in the presence of different IgM concentrations: **(A)** Probability density function (PDF) histogram represents unfolding time of protein L in the absence of IgM (control) is ∼1.9 s. **(B)** In the presence of 1µM IgM, the peak of unfolding time was increased to ∼33.6 s. **(C)** This unfolding time further upshifted to ∼66 s with 6 µM IgM. **(D)** These peak unfolding times have been plotted against the IgM concentrations with double exponential fitting, indicating two plausible binding sites with different avidity. The analysis ha been performed by bootstrapping the data (N=100) from 7 different molecules without IgM (control, A), 8 molecules with 1 µM (B), and 7 in the presence of 6 µM IgM (C).

**Figure 3:**
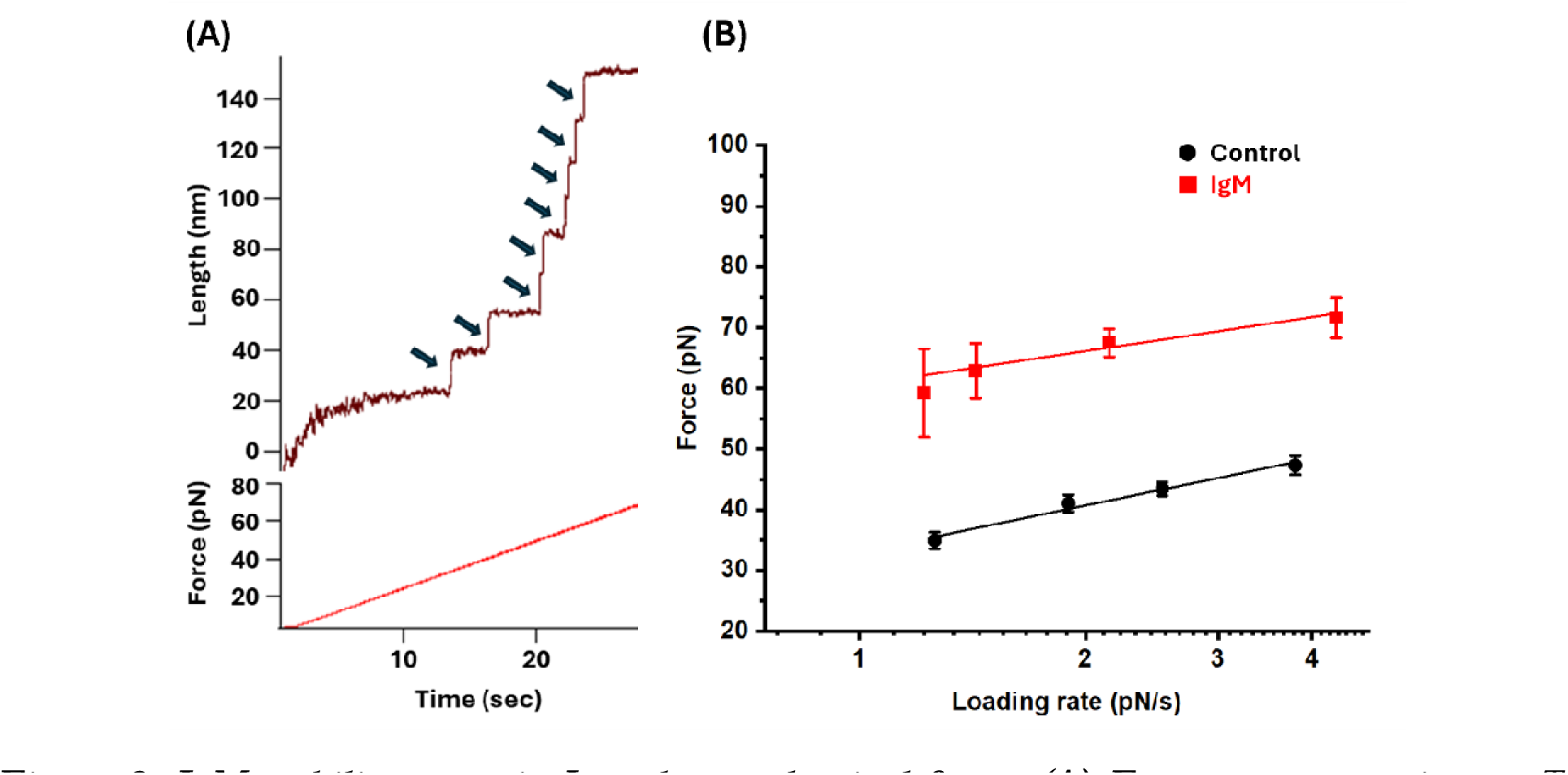
IgM stabilizes protein L under mechanical force: **(A) Force-ramp experiment:** To assess the mechanical robustness of the octameric protein L construct, force was gradually increased from 4 to 80 pN via a constant increasing ramp rate of 2.53 pN/s, followed by decreasing ramp at the same rate. Unfolding force was determined during the force-increase ramp by extrapolating the unfolding steps to the force-ramp curve. **(B) Unfolding force of Protein L at different force-ramp rates:** Unfolding forces of protein L were calculated at various ramp rates, within 1.2-4.3 pN/s range. The mechanical stability of protein L increased in the presence of IgM, as evidenced by an increase in unfolding forces from 32.98±0.93 pN to 60.54±1.63 pN at a zero-ramp rate. Data points represent measurements from more than three individual molecules at each ramp rate, with error bars indicating standard error (s.e.).

Since this induced mechanical stability of protein L is rendered by their structural rearrangement upon IgM binding, it depends on the response of their antibody-binding interfaces that be observed as their distinct conformational trajectories during mechanical unfolding of the protein L-IgM complex. To investigate the contribution of these binding interfaces on the elevated mechanical stability of protein L, we have further performed simulation study of Protein L-IgM complex to explore the dynamic nature of their intermolecular interactions under mechanical constraints.

### The molecular basis of protein L-IgM interaction revealed by steered molecular dynamics (SMD) simulations

Protein L B1 domain (PDB 1HZ6) has a simple topology with a single α-helix that is packed against four stranded β-sheets (Supp. Fig. 1). To prepare suitable model structures of the Protein L B1 domain bound with one or more IgM antibodies, we need to identify the interacting residues between these two proteins at the binding interface. We used this information from an earlier study by Graille et al. and then performed site-specific docking using the HDOCK platform^9,16^ (Supp. Table 1). These model structures of the complex were used to carry out atomistic molecular dynamics (MD) simulations for 200 ns each for three systems: (i) apo state, (ii) two monomer systems: IgM bound to site 1 of Protein L (binding via C-terminal β-sheet with α-helix) and site 2 (binding via N-terminal larger β-sheet) separately (Fig. 4), and (iii) dimer system: two IgM antibodies bound to both site 1 and site2 (Fig. 5). The stability of these modelled structures in MD simulations was validated using standard structural parameters: root-mean square deviation (RMSD), solvent-accessible surface area (SASA), radius of gyration (RoG), and the distance between two Cα pulling atoms (Supp. Fig. 2; Supp. Table 2). No significant perturbation has been observed in these complexes, suggesting that the protein L-IgM complexes are highly stable.

**Figure 4:**
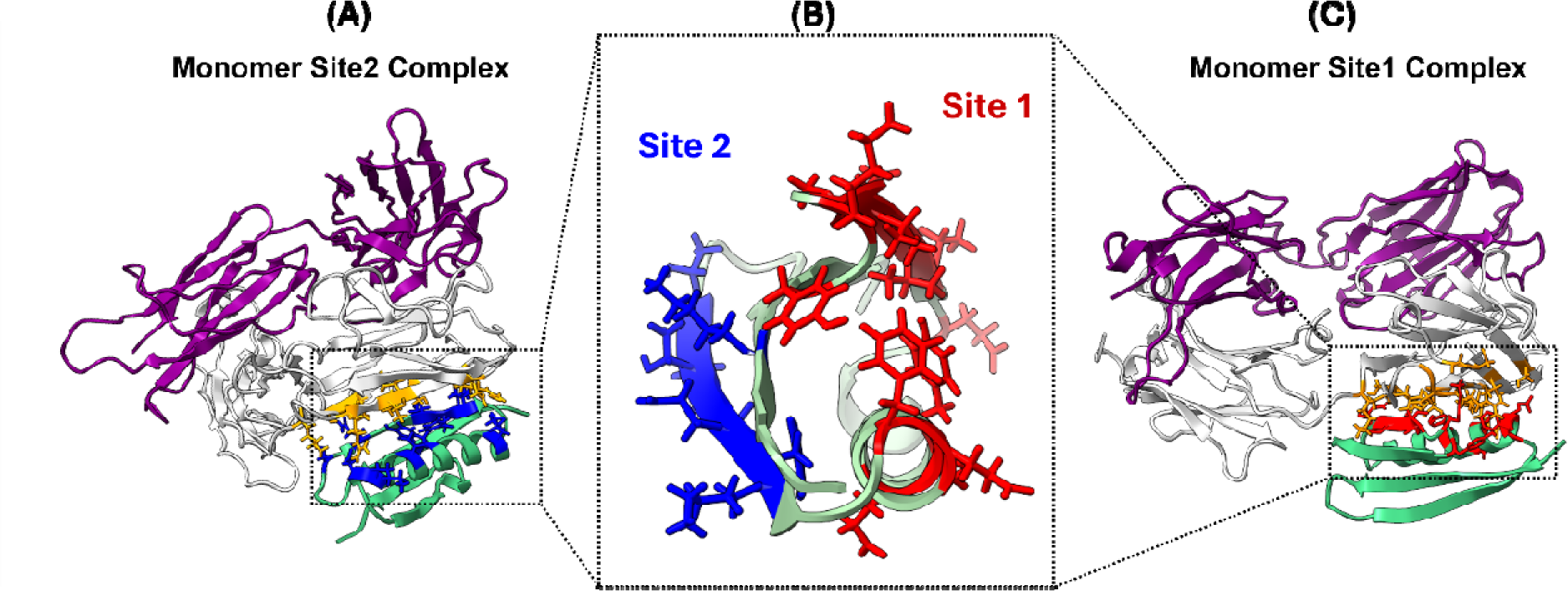
Protein L-IgM monomer complex formation: (A and C) Binding interfaces of protein L and IgM : Protein L (sea-green) interacts with the chain of IgM molecule (white) either through the site 1 or site 2. In both the cases, the IgM interacting residues are visualized as orange sticks. The interacting residues of protein L site 1 are indicated by red sticks, while site 2 by blue sticks **(B)**.

**Figure 5:**
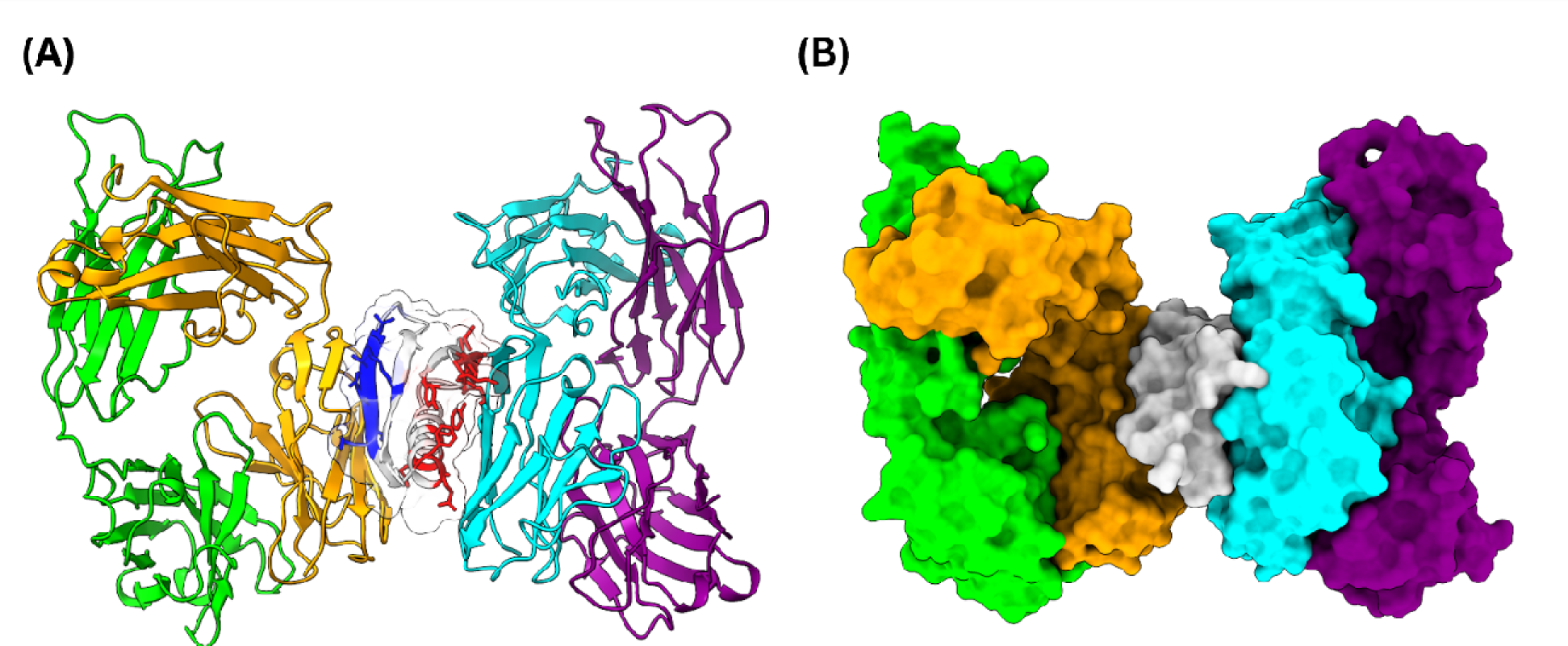
Protein L complex formation with two IgM molecules: (A) Binding interface during dimer complex formation: During the dimer complex formation, protein L simultaneously binds to two IgM molecules, through both their site 1 and site 2, which are indicated by red and blue sticks, respectively. **(B) Surface volume occupancy:** Surface volume occupancy has been depicted of protein L (grey) with 2 IgM antibodies through both site 1(right, IgM chain i indicated by cyan) or through site 2 (left, IgM chain is indicated by orange).

To prepare the dimer protein L-IgM complex, we initially used the replacement method-first superimposing protein L structure from both PDBs 1HEZ and 1HZ6 in the same orientation with IgM bound state, and then removing the 1HEZ to create a complex between IgM and 1HZ6 through both site1 and site 2. Subsequently, we have checked their stability through equilibrium MD simulation as other complexes.

For all the IgM bound Protein L systems, we have calculated the binding energy between Protein L and IgM using the MM/GBSA method over the last 100 ns of each equilibrium MD trajectory (Supp. Table 3). We must clarify that MM/GBSA method has several limitations and should not be trusted for absolute binding free energy calculations. However, here our goal is to establish a qualitative trend of the binding energies. As observed from Supp. Table 3, the site 1 has stronger affinity compared to site 2. Thus, it is expected that at lower antibody concentrations, site 1 will be occupied preferentially. With increasing antibody concentration, the dimer population will start to increase. Hence, the experimentally observed forced profile is likely to be observed as a weighted average of a certain population of the monomer and site 1, monomer at site 2 at lower concentrations and primarily dimer at higher concentrations.

We have performed steered molecular dynamics (SMD) simulations starting from the well-equilibrated protein L-IgM complexes and the apo protein L systems to calculate their unfolding rupture force profile. SMD pulling was done at a constant velocity of 0.005 Å/ps. The extension with time curves has been shown for all the systems (Supp. Fig. 3). Interestingly, we have observed that the rupture force increases steadily from 1773.97±46.97 pN for apo-protein to 8991.21±65.88 pN in the dimer complex (Fig. 6, Supp. Table 4). Since the binding affinity of protein L and IgM is directly attributed to the work done (kcal/mol) profile, we observed a similar trend in work done among these complexes (Fig. 8).

**Figure 6:**
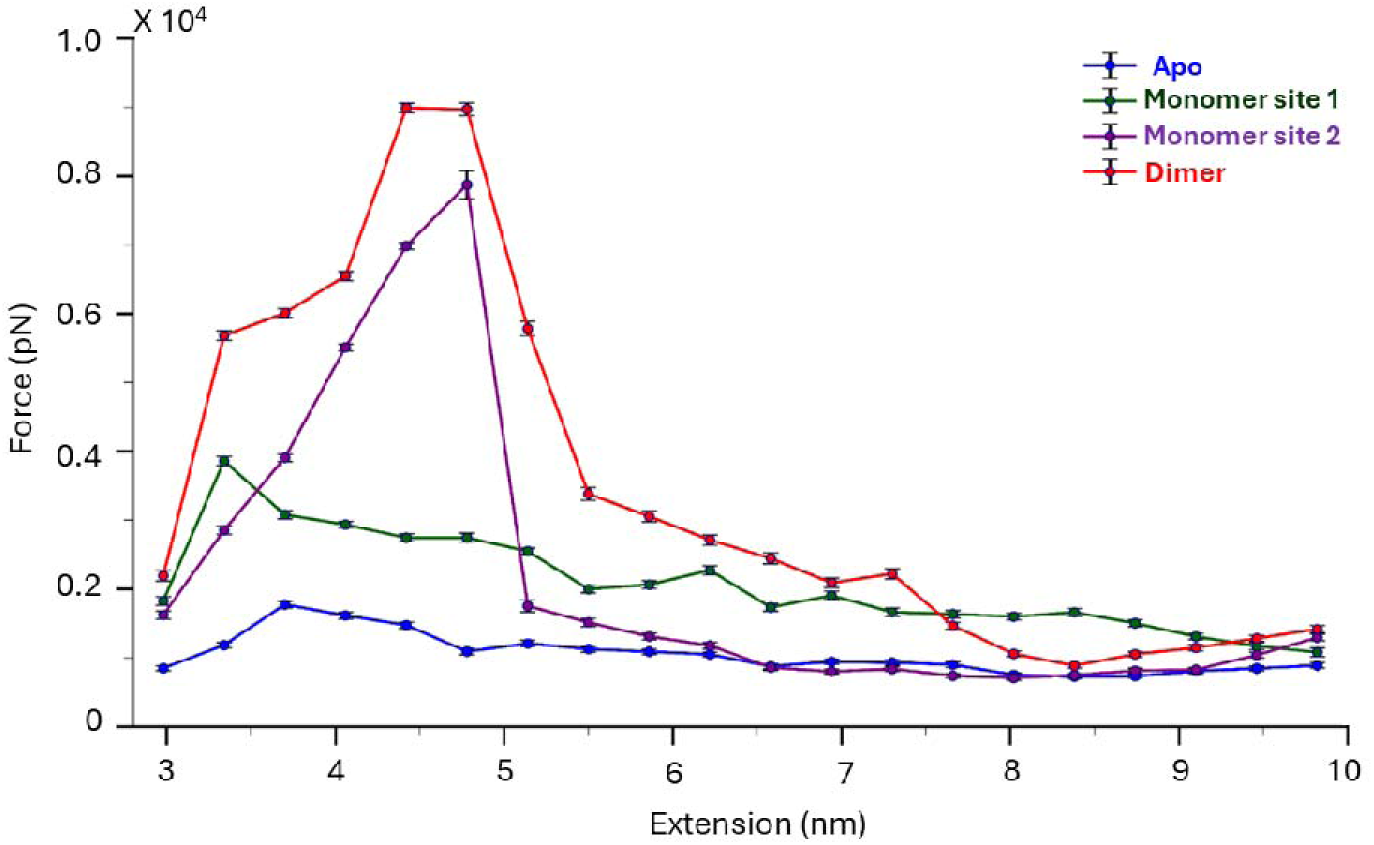
Rupture pulling force profile of protein L-IgM complex: The Protein L-IgM complex has been pulled at a constant rate of 0.005 Å/ps to determine the rupture force of protein L complex and plotted against the extension trajectory. The force profile has been determined for four different conditions: apo state (indigo), monomer site 1 (complex with single IgM via site 1, green), monomer site 2 (complex with single IgM via site 2, purple), and dimer (complex with two IgM antibodies through both site 1 and site 2, red). We have observed that the rupture force value increases steadily from 1773.97±46.97 pN in case of apo-state to 8991.21 ±65.88 pN in dimer complex formation, suggesting the IgM-induced mechanical stability of protein L. The pulling experiments have been performed three times and plotted as average with standard error for three runs.

To explain the underlying molecular principle behind the IgM-induced stability of protein L, we dissected the sequence of unfolding events for protein L in the apo state and different IgM complexes (site 1, site 2 and both). We observed that in the absence of any IgM molecule, the initial stages of unfolding of protein L involve the breaking of its intrachain contacts between two parallel β-sheets, which triggers the unfolding of N-terminal β-sheets (site 2), followed by the loosening of intrachain contacts of C-terminal β-sheets with α-helix (site 1). Eventually, within 10 ns of the pulling run, the β-sheets completely unfold, and the entire structure was fully dismantled within 20 ns of the SMD run (Fig. 7, 1^st^ row).

**Figure 7:**
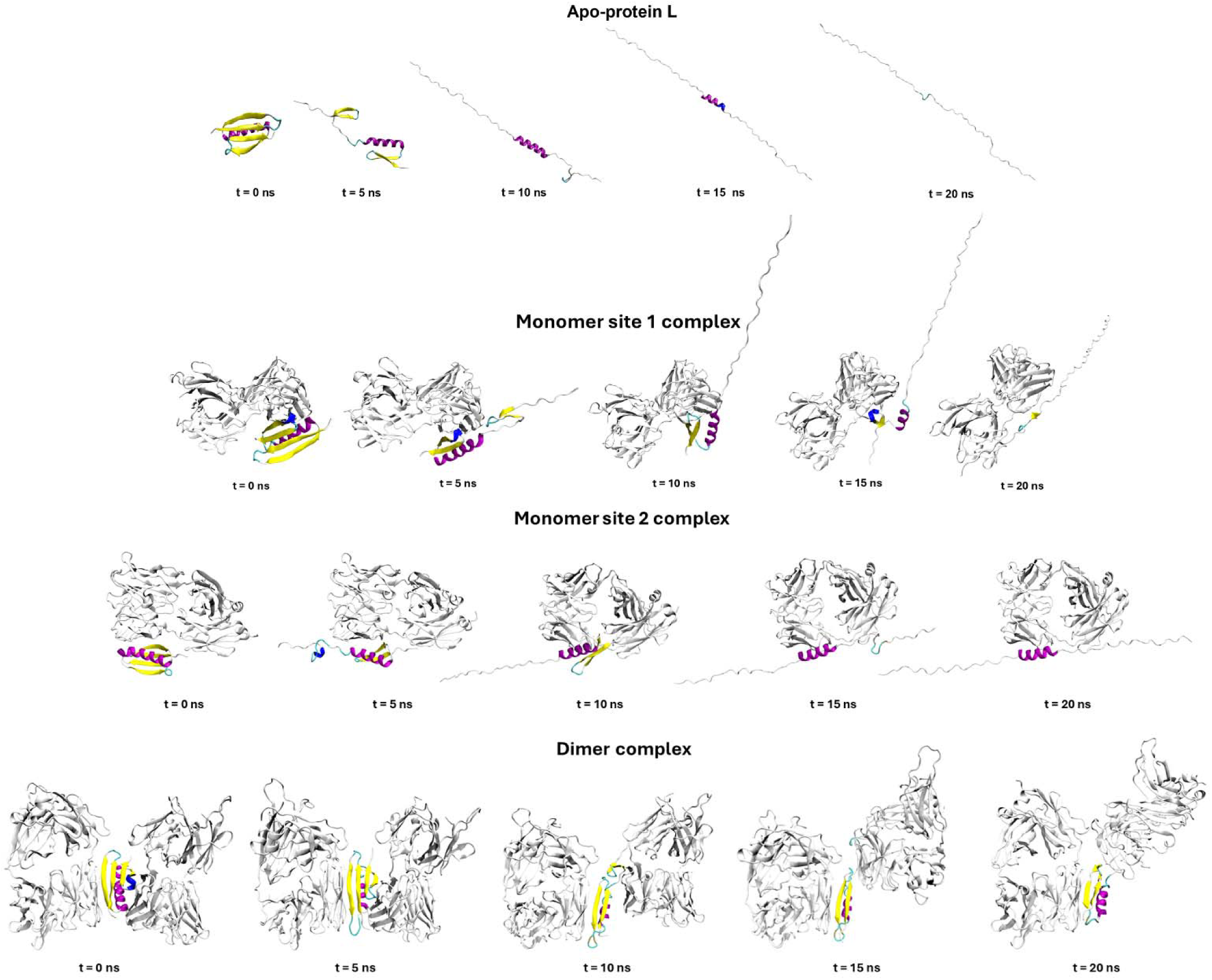
Molecular events of protein L unfolding at different pulling timescales: Apo-protein L (1^st^ row): In case of apo-protein L, there is an initial breaking of intrachain contacts between the two parallel beta sheets within the structure, leading to the unfolding of N-terminal β-sheet. Following it, the C-terminal smaller β-sheet loses its intrachain contacts with the α-helix, and eventually unfolds. At last, the remaining α-helix got disrupted and the entire structure was fully dismantled within 20 ns of the SMD run. **Monomer site 1 complex (2^nd^ row):** In case of monomer site 1 complex, the C-terminal β-sheet and α-helix are responsible for making interchain interactions with the IgM chains. During the pulling exercise, the non-interacting N-terminal β-sheet was unfolded and completely lost its structure within first 7 ns of run. Despite this unfolding, the interacting part of the protein L was found to retain its native scaffold up to 12.5 ns of the simulation run. At ∼15 ns, the C-terminal β-sheet stars losing its secondary structure, followed by α-helix. Interestingly, it was observed that the smaller significant portion of the C-terminal β-sheet retained its secondary form and was in close contact with the IgM chain. Further, bond breaking frequency got delayed by the presence of the antibody, thus improving the mechanical stability. **Monomer site 2 complex (3^rd^ row):** By contrast to monome site 1 complex, monomer site 2 complex undergoes the unfolding from its C-terminal β-sheets and associated α-helix segment, while N-terminal parts retain folded scaffold due to their interchain interaction with IgM. However, the unfolding response is slower than the monomer site 1 complex, suggesting a higher mechanical stability. **Dimer complex (4^th^ row):** During the pulling of the dimer complex with IgM, we found that site 1 unfolds quickly in the direction of applied force. However, the site 2 was found to be in complete folded form under strong contact with the IgM chain throughout the simulation run. In addition, the applied force was not found to be enough for disrupting the site 2. This suggests that the site 2 interface possesses higher mechanical stability than the site 1 upon IgM interaction; thus, the overall mechanical stability of dimer complex was found to be higher.

By contrast, in the case of the monomer site 1 complex, the C-terminal β-sheet and α-helix are responsible for making interchain interactions with the IgM chains. In the SMD pulling runs, the non-interacting N-terminal β-sheet was unfolded first and completely lost its structure within the first 7 ns. Despite this unfolding, the interacting part of the protein L was still found to retain its native scaffold almost up to 12.5 ns. Interestingly, the smaller significant portion of the C-terminal β-sheet retained its secondary structure and was in close contact with the IgM chain. As a result, the pulling forces were not able to break this secondary structure of this segment due to the presence of the antibody (Fig. 7, 2^nd^ row).

For the system with IgM bound to site 2, N-terminal β-sheet and α-helix of protein L engage with the IgM chains. The non-interacting C-terminal β-sheets are more susceptible to unfold under mechanical constraints within 5ns run. However, eventually, the N-terminal β-sheet was observed to initiate unfolding and losing its structure via breaking the intrachain contacts. Intriguingly, throughout the pulling run, the interacting α-helix with the IgM chain was found to be in the folded conformation. The inability of the applied force to completely disrupt the structure is consistent with the observed higher peaks in the rupture force profile, highlighting comparatively higher mechanical stability of protein L-IgM complex formed through site 2 than site 1 (Fig. 7, 3^rd^ row). In the dimer complex, where both site 1 and site 2 are engaged with IgM, we observed that site 1 unfolds first under pulling forces. However, the site 2 interface was found to retain the structural integrity through forming a strong association with IgM (Fig. 7, 4^th^ row).

## Discussions

Protein L is a well-known super-antigen located on the bacterial surface that interacts with human antibodies. These antigenic proteins have evolved to function under and withstand shear stress generated by diverse physiological processes such as mucus flow or coughing^5,17,18^. Our single-molecule results have provided an empirical demonstration that IgM association augments the mechanical stability of protein L, and this effect could be regulated by the two IgM binding interfaces on protein L. This empirical evidence elucidates the intricate molecular mechanisms governing the protein L-IgM interaction, elucidating their role in ensuring the stability and functionality of the protein amidst dynamic physiological conditions.

It is already known that protein L has two binding sites with different avidity; however, their individual contributions to the mechanical strength of protein L are not well understood^9,19–21^. When the antibody concentration is low in the substratum site, the high-avidity binding interface of protein L is more likely to act; however, their second binding site with low-avidity aids in promoting the overall mechanical stability of the protein L during the shear flow at high antibody concentration. Interestingly, we found that with the IgM molecule, protein L site 2 is the high-avidity site, which shows higher mechanical stability than with site 1, i.e. low-avidity force-sensor site (monomer site 1 vs monomer site 2 complex, Supp. Table 4). Interestingly, we observed that IgM association to both these sites (i.e. dimer complex) shows an additive effect on the mechanical stability of the protein L-IgM complex, promoting the highest rupture force (Supp. Table 4). This synergistic stabilizing effect of both the binding interfaces is responsible for delaying the unfolding time of protein L, that has been demonstrated by single-molecule pulling experiments (Fig. 2). Therefore, it can be assumed that at low IgM concentration, protein L already gains the mechanical stability through the site 2 interface. Since site 2 acts as the larger engagement tether during primary IgM exposure, it individually promotes higher mechanical stability and during this tethered-scanning, protein L plausibly extends its domain in such a way that it increases the search area and connects with another antibody through the site 1. This further IgM binding through the site 1, as a secondary one, only supports in increasing the mechanical stability of the protein L complex to withstand the immune response under force. More importantly, in case of dimer complex, low-avidity site 1 of protein L unfolds prior to the high-avidity site 2, corroborating their role as initial force-sensor. It also suggests the higher mechanical stability of site 2 than site 1 upon IgM association under physiological scenario.

Based on our data, it is reasonable to anticipate that the high-avidity site 2 is more mechanically resilient, that enables bacteria to ascertain substrate attachment even in the presence of low antibody concentration. Once attached on the substrate, the local antibody concentration increases and the IgM-induced mechanical stability of site 1 promotes the structural rigidity to protein L, producing a mechanical signal to inform about the antibody concentration at infection site. Future investigations may involve single-molecule experiments utilizing protein L mutants with alterations in their binding interface residues. This research avenue aims to elucidate the systematic impact of these binding sites on protein L stability upon antibody interaction. Overall, this mechanosensitivity of protein L domain through its two binding sites can illustrate their regulated mechanical stability in the presence of varying concentration of these different antibodies at different phases of infection.

## Materials and methods

### Expression and purification of protein L

The protein L polyprotein construct has been transformed into *E. coli* BL21 (DE3) cells and then were grown in the Luria broth with Carbenicillin (50 µg/ml) at 37° C until the O.D._600_ reaches 0.6-0.8, followed by induction with 1 mM Isopropyl β-D-thiogalactopyranoside (IPTG, Sigma Aldrich) for overnight at 25°C. On following day, the cells were pellet down by centrifuging at 9000 rpm and then resuspending in 50 mM sodium phosphate buffer 300 mM NaCl and 10 % glycerol at pH 7.4. The dissolved cell pellet was incubated with phenylmethylsulphonyl fluoride and lysozyme at 4°C for 20 minutes and was further treated with DNase, RNase, Triton-X 100, and MgCl_2_ and kept on the rocking platform at 4°C. For cell disruption, mechanical cell homogenizer (Constant systems) was used at 19 p.s.i., followed by centrifugation at 4°C. The protein was purified from the supernatant collected, with Ni^2+^-NTA column chromatography of AKTA Pure (GE Healthcare) with the binding buffer, comprising of 20 mM imidazole, 50 mM sodium phosphate and 300 mM NaCl with pH 7.4. Same buffer composition was used for elusion with 250 mM imidazole. Next, the protein was concentrated by Amicon concentrator and biotinylated as per manufacturer protocol (Avidity). Briefly, the reaction mix was kept at 30°C with intermittent shaking for 2 h and then kept at 4°C for overnight. On the next day, size exclusion chromatography was performed by the Superdex-200 increase 10/300 GL.

### Glass chamber preparation

The bottom glass slides were cleaned by sonicating for 15 min with Hellmanex III (1.5%) solution at 60°C followed by washing them with double distilled water. Cleaned cover slides were then treated with a mixture containing methanol (CH_3_OH) and concentrated hydrochloride acid (HCl), followed by concentrated sulphuric acid (H_2_SO_4_). To activate glass slides through silanization, the slides were suspended in the ethanol solution containing 1% (3-Aminopropyl)-trimethoxysilane for 15 minutes and washed gently with ethanol for removing the unreacted silane from the chamber and dried for 1 hour. Similarly, coverslips were cleaned by washing them with Hellmanex III (1.5%) solution and then treated with ethanol^22^. By sandwiching the glass slides and the coverslips together with parafilm, a chamber is prepared. Then glutaraldehyde (Sigma Aldrich) solution is passed through the chamber and left 50 min to react followed by reference beads (2.5-2.9 µM, Spherotech, AP-25-10) and O4 ligand (Promega, P6741). At last glass chambers were flushed with blocking buffer (150 mM NaCl, 20 mM tris-HCl, 2 mM MgCl2, 0.03% NaN_3_ and 1% bovine serum albumin, pH 7.4) and left for 5 hours at room temperature to avoid any nonspecific interactions.

### Single-molecule magnetic tweezers experiments

A bespoke magnetic tweezers was crafted, utilizing a Zeiss Axiovert S100 microscope as the foundational platform to carry out single-molecule experiments, as described previously^7^. The microscope stage hosts a glass flow chamber, which is illuminated by a collimated cold white LED sourced from Thor Labs. The 63X oil-immersion objective is used to display beads visuals, and the P-725 nanofocusing piezo actuator is also attached. Ximea camera (MQ013MG-ON) is used to process Images. A multifunctional DAQ (Ni-USB-6289, National Instrument) is used to control both the piezo positioning and data acquisition. Magnetic force is applied using neodymium magnets attached to the linear voice coil (Equipment Solutions). The force’s intensity can be adjusted by varying the vertical position of the magnets along the z-axis. Detailed methods for force calibration, bead tracking, and image processing have been previously outlined^12,22^. Experiments took place within a flow chamber containing protein concentrations ranging from 1 to 10 nM in 1X PBS buffer (pH 7.2). We obtained the antibodies through commercial suppliers: IgA from human serum (Sigma-Aldrich), IgM from human serum (Sigma-Aldrich). We prepared the antibody stock solution in 1X PBS (pH 7.2) and used a range of concentrations. The range of concentration 0-6 µM for IgM; and 0-6 µM of IgA were used in the single-molecule experiments.

### Structures of Protein L protein and multiple sequence alignment (MSA) studies

From the Protein Data Bank database^23^, 3 initial structural coordinates of substrate protein L was fetched; PDB entries 1ZH6 (Resolution: 1.70 Å), 1MHH (Resolution: 2.10 Å), 1HEZ (Resolution: 2.70 Å) (Supp. Table 2). In this study, PDB entry sequence 1ZH6 is the protein of interest which was used for experimental studies and the PDB entry 1HEZ was taken as a reference template (dimer complex).

From the respective PDB entries, the Protein L sequence was pulled out and used for multiple sequence alignment using the Multalin platform^24^ (Supp. Fig. 4). The alignment studies confirmed the variations in the sequences of protein L with respect to PDB 1ZH6.

### Molecular docking and complex generation studies

Docking studies for the substrate protein L (1ZH6) and the target receptor (IgM antibody) were executed via the online HDOCK platform^16^. This platform utilizes a hybrid docking algorithm coupled with a fast Fourier transform (FFT)-based global search and an iterative knowledge-based scoring function. Each complex yielded 100 docked poses, meticulously evaluated to identify the optimal pose by considering the scores of their interacting residues and binding energies. The 3D docked complexes for both sites and the residue level interactions (within 4.5 Å) was visualized using ChimeraX platform^25^ and noted down (Supp. Table 1.). Additionally, the 2D interaction plots were also generated using the DIMPLOT module of LIGPLOT software (Supp. Fig. 5 and 6).

For the dimer complex, we had employed the replacement method, in which we superimposed the protein L structures from PDBs 1HEZ and 1HZ6 using Chimera X matchmaker module, such that both structures were oriented similarly to the IgM-bound state. Afterward, we excluded the structure from 1HEZ (Protein L section), which led to the formation of a dimer complex between IgM and 1HZ6, engaging both site 1 and site 2.

### Molecular dynamics studies

To understand the stability of generated complexes, MD simulations were carried out using GROMACS 2019.6 software package^26^. The protein parameters for the simulation runs were created using the CHARMM-36 all-atom force field^27^. Prior to the production run, the complex systems were enclosed within a cubic box filled with the TIP3P water model and simulated with periodic boundary conditions^28^. The residues were adjusted according to their standard ionization states at pH 7.0. The entire complex was neutralized with a 0.15 M NaCl solution to replicate experimental salt conditions. The simulation was carried out in three parts, using a force constant of 1000 kJ/mol. nm² to restrain all the heavy atoms and maintain the original protein folding. The first part involved the initial optimization of each system’s geometry for 5 ps using the steepest descent algorithm. This was followed by a two-stage equilibrium process, with 100,000 iterations (100 ps) at each stage. The initial equilibration was performed through a constant number of particles, volume, and temperature (NVT) ensemble using the Nose-Hoover temperature coupling method for controlling the temperature within the box^29^. The subsequent equilibration was conducted under constant under a constant pressure and temperature (NPT) ensemble at 1 atm and 303.15K using the Parrinello-Rahman barostat^29^. During the simulation, long-range electrostatic interactions were calculated using the particle mesh Ewald (PME) summation method^30^. Short-range electrostatic and van der Waals interactions were cutoff at 1.2 nm. Throughout the production run, covalent bonds were constrained using the LINCS algorithm^31^, and a time step and collection interval of 2 fs and10 ps were set. The systems of interest underwent a 200 ns dynamics production run under constant pressure conditions.

Additionally, the simulation trajectories were analyzed to assess the root mean square deviation (RMSD), radius of gyration (ROG), solvent-accessible surface area (SASA) values using gmx RMSD, gmx RMSF and gmx gyrate tools, respectively^26^ to comment on the stability of the complexes under dynamics environment.

### Steered molecular dynamic (SMD) simulation

For driving the molecular systems to an unfolding state, and capturing the mechanical stability of the system, a constant velocity Steered Molecular Dynamics (SMD) simulation exercise was performed using GROMACS 2019.6 software package^26^ patched with PLUMED version 2.7.4^32^. Initially, before the pulling exercise, the system was prepared and well equilibrated under standard equilibrium Molecular Dynamics (MD) conditions. In this study, end-to-end CA atoms (N-terminal Glu residue as Cα atom, and C-terminal Gly residue as Cα atom) pulling were carried out for the PRO L across all the systems (Supp. Fig. 7). In addition, the mean interatomic spacing between the pulled atoms of the equilibrated complexes (before SMD), was calculated and tabulated down (Supp. Table 2).

The constant velocity pulling was conducted until the end-to-end distance reached 10 nm from the initial CA distance, and it was performed over a duration of 20 ns, at a rate of 0.005 Å/ps and a spring constant (κ) of 1000 kJ/mol/Å^2^, this exercise was repeated for 3 iterations runs for each system. The pulling trajectory, rupture force, the extension distance, the work done datapoints, were recorded in the Colvars configuration file at intervals of every 5000 steps. The COLVAR pulling datasets was further used for calculating and constructing the rupture force profile along with standard errors (Supp. Table 4) and visualization in Fig. 6. The amount of work done during the rupturing exercise of each complex was also constructed and visualized (Fig. 8), incorporating data from three levels of iterations run (Supp. Fig. 8-13).

**Figure 8:**
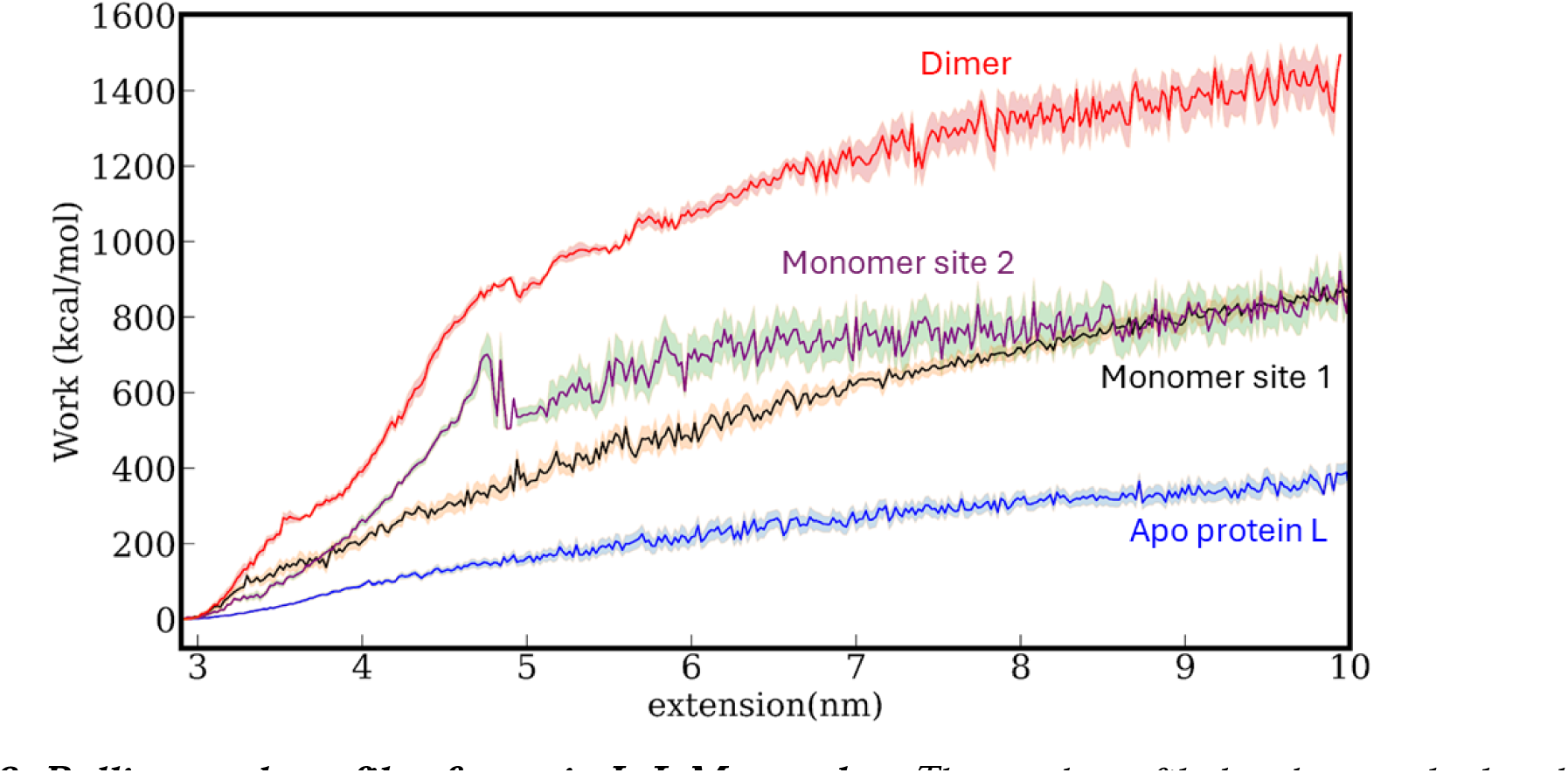
Pulling work profile of protein L-IgM complex: The work profile has been calculated from the simulated pulling trajectory of the respective complexes. We found that the work done increases significantly, complementing the force profile; and becomes almost ∼5 times in case of dimer complex. Simulation at each condition has been run for three times and the shaded regions indicate the standard error for each condition.

The extracted trajectories of the complexes were utilized for the purpose of analyzing the protein L unfolding crucial molecular events over the course of dynamic run using VMD (visual molecular dynamics) tool^33^, and the time frame evolution of the unfolding mechanism across the complexes were also captured (Fig. 7).

### MM-GBSA based free energy calculation studies

To provide the thermodynamic essence, the free energy of binding of substrate protein L with the receptor chain of interest was calculated using a MD based-Molecular mechanics with generalized Born and surface area solvation (MM/GBSA) method^34^. GMX_MMPBSA tool^35^ helps us to estimate ligand-binding affinities against a receptor choice of interest. A total of last 1000 frames (100 ns frames) from a well equilibrated stable 200 ns trajectory at an interval of 1 frames difference were extracted and used for the calculation of the mean total binding energy of the system in complex formation (Supp. Table 3), the entropy contribution was not taken into consideration for determining the ligand binding free energy (BFE) calculation. The overall energy of the system was determined using the equation: ΔG_Binding_ = ΔG_Complex_ - ΔG_Receptor_ - ΔG_Inhibitor_.

## Author Contributions

S.C.^1^, S.H.^1,2^, and S.C^2^. designed the project. S.C.^1^ and M.B.^1^ performed the experiment. S.C.^1^ analyzed the experimental data. S.P.^2^ carried out the computational work and data analysis. S.C.^1^, S.H.^1,2^, S.P.^2^ and S.C.^2^, D.C.^2^, wrote the manuscript.

## Acknowledgemen**t:**

We thank S.N. Bose National Centre for Basic Sciences, Technical Research Centre (TRC) project of Department of Science and Technology (DST), Ashoka University and the Mphasis foundation for support and funding. S.H. thanks DST SERB Core Research Grant for funding. S.C^2^ thanks SERB, DST, India for funding.

